# Impact of Water Deficit on Growth, Biochemical, and Physiological Traits in Eggplant MAGIC Lines

**DOI:** 10.64898/2026.09.01.748492

**Authors:** Martín Flores-Saavedra, Mariola Plazas, Nuria Pascual-Seva, Santiago Vilanova, Oscar Vicente, Pietro Gramazio, Jaime Prohens

**Author notes:** Corresponding author: E-mail address (M. Flores-Saavedra).

## Abstract

Climate change exacerbates agricultural water scarcity, necessitating the development of drought-tolerant crop varieties. This study evaluates 12 eggplant lines from a MAGIC (Multi-parent Advanced Generation Intercross) population, previously selected for contrasting responses to water deficit during the vegetative stage. To validate tolerance under adult production conditions, plants underwent five irrigation-withholding cycles over a 170-day greenhouse growing period. Yield components, the Stress Tolerance Index (STI), and physiological parameters (water status and stomatal conductance) were evaluated. Additionally, photosynthetic pigments, oxidative stress markers, antioxidant compounds, and osmolytes were quantified to characterize the biochemical basis of tolerance alongside final biomass production. The results showed that four of the five lines that were previously classified as tolerant in the vegetative stage remained among the most tolerant at the reproductive stage. Specifically, lines L13, L78 and L179 were the most productive under water-limited conditions. While L13 and L179 exhibited stable tolerance throughout all developmental stages, L78 displayed stage-specific tolerance, manifested only during the reproductive growth phase. These findings emphasise the importance of integrating early-stage screening with adult-stage validation in order to capture the full spectrum of genetic drought tolerance. The most productive lines were characterised by moderate aboveground biomass, high leaf hydration and maintained stomatal conductance. However, the strategies employed differed: while L179 exhibited high photosynthetic pigment content, L13 was characterised by high total sugar accumulation. Overall, these results provide a multi-trait roadmap and identify elite MAGIC parental lines for breeding climate-resilient eggplant cultivars.

## 1. Introduction

Changes in weather patterns pose a threat to food security, causing significant losses in agricultural productivity (Hultgren et al., 2025). Rising temperatures have intensified evaporative demand, contributing to greater aridity in many regions. Furthermore, population growth has increased the demand of water for agriculture, a resource that is becoming increasingly scarce (Palmgren & Shabala, 2024). Climate change is progressively affecting water availability for irrigation through changes in rainfall patterns and increased crop evapotranspiration. Therefore, it is essential to improve water use efficiency through agricultural practices and the use of drought-resilient crops (Muzammal et al., 2024).

Agriculture is the largest user of water worldwide. Horticultural crops, in particular, require a considerable amount of water, especially in arid or semi-arid regions (Ferreira et al., 2024). Drought stress is one of the main factors limiting horticultural production, affecting quality and increasing the incidence of pests (Bisbis et al., 2019). Amongst horticultural crops, eggplant (*Solanum melongena* L.) is widely cultivated in regions where water scarcity is becoming a growingly constraint (Toppino et al., 2022), making this crop especially relevant in the context of intensifying water deficit. In this way, drought negatively impacts eggplant growth, resulting in reduced gains in both aerial and root biomass, as well as smaller leaf area and plant height (Flores-Saavedra et al., 2023). Eggplant production is significantly affected by reduced irrigation. Specifically, yields have been shown to decrease by up to 39% when plants receive only 50% of their water requirements compared to well-irrigated conditions (Maachi et al., 2025). However, eggplant can tolerate mild stress conditions without significantly compromising yield (Díaz-Pérez & Eaton, 2015). Reduced water availability also impacts the quality of eggplant fruit, resulting in smaller fruits with a higher soluble solids content (Çolak et al., 2017).

Under water deficit conditions, plants undergo a series of physiological and biochemical changes, including stomatal closure, reduced gas exchange, increased production of reactive oxygen species, accumulation of osmolytes and altered responses to phytohormones. These changes affect plant growth and development (Khalid et al., 2023). Eggplant plants exhibit a greater reduction in photosynthesis and stomatal conductance when water availability is low (Díaz-Pérez & Eaton, 2015; Tani et al., 2018). Photosynthetic pigments in eggplant leaves are also negatively affected by water deficit, with chlorophyll and carotenoid content decreasing under drought conditions (Plazas et al., 2019). Furthermore, drought induces oxidative stress in eggplant, increasing H₂O₂ and MDA content. In this context, phenolic compounds have been observed to play a key an antioxidant role (Flores-Saavedra et al., 2024; Plazas et al., 2019). Regarding the cellular accumulation of osmolytes in plant, proline content has been extensively studied under different stress conditions. In eggplant, proline is a fundamental secondary metabolite that helps the plant make physiological adjustments in response to stress, for example to drought treatments (Chakhchar et al., 2025).

Screening plant genetic resources is essential for identifying genotypes that are tolerant of water stress. These genotypes adapt through a strategic combination of physiological, morphological, and biochemical traits (Kumar et al., 2012). The large genetic diversity of the eggplant, alongside its close phylogenetic relationship with over 500 *Solanum* species, represents a strategic opportunity to improve adaptive traits in response to abiotic stress (Gramazio et al., 2023). These resources include experimental populations that incorporate wild species among the founder parents, which are particularly noteworthy. Examples include introgression lines of *S. incanum* with favourable alleles for root growth and chlorophyll content in water deficit conditions (Flores-Saavedra, et al., 2025), or an eggplant multiparent advanced generation intercross (MAGIC) population that includes the eggplant wild relative *S. incanum* as one of its parents (Arrones et al., 2025). This MAGIC population has enabled the selection of lines with greater tolerance and the identification of genomic regions associated with drought-tolerant traits in young plants (Flores-Saavedra, et al., 2026).

This study aims to evaluate the impact of water deficit on yield, growth, and the physiological and biochemical profiles in a subset of 12 eggplant MAGIC lines. Furthermore, we seek to identify genotypes with superior drought tolerance and to pinpoint the specific traits that underpin resilience and optimal development under water-limited conditions.

## 2. Materials and Methods

### 2.1. Plant material

The study evaluated 12 lines from a MAGIC population of eggplant, comprising seven cultivated eggplant (*S. melongena*) parents and one parent corresponding to the wild relative, *S. incanum* (Arrones et al., 2025). These 12 lines were selected due to their contrasting responses to stress during the vegetative phase in a previous population trial (Flores-Saavedra et al., 2026). This trial examined traits associated with greater drought tolerance, such as water content, total growth, root growth, and proline content. Six of the lines were classified as tolerant (L13, L38, L56, L59, L179, and L323), and six as susceptible (L57, L78, L101, L234, L242, and L277) (Flores-Saavedra et al., 2026).

### 2.2. Growing conditions

The seeds of the 12 lines were germinated in Petri dishes according to the germination protocol described by Ranil et al. (2015). After germination, the seedlings were transferred to seedling trays containing a growing substrate (Humin Substrate N3, Klasmann-Deilmann, Geeste, Germany) in a growth chamber. On 10 June 2025, forty-one days after germination, the seedlings were finally transplanted into 15 L pots containing coconut fibre (Horticoco, Valimex, Valencia, Spain) in a benched Venlo-type greenhouse located at the Universitat Politècnica de València (Valencia, Spain). The pots were arranged in a randomised complete block design with five blocks, with each line represented by one plant per block in each treatment. Within each bench, pots were spaced 0.6 m apart, while 1.2 m was maintained between benches. The plants were manually pruned to control lateral growth and trained to two stems per plant. Greenhouse climate conditions were monitored throughout the experimental period (June to November 2025) using an ATMOS14 sensor (METER Group, Pullman, WA, USA). During this period, the average maximum and minimum temperatures were 30.1°C and 21.9°C, respectively, whereas the average maximum and minimum vapour pressure deficits were 2.17 kPa and 0.58 kPa, respectively (Figure 1a, b).

**Figure 1.**
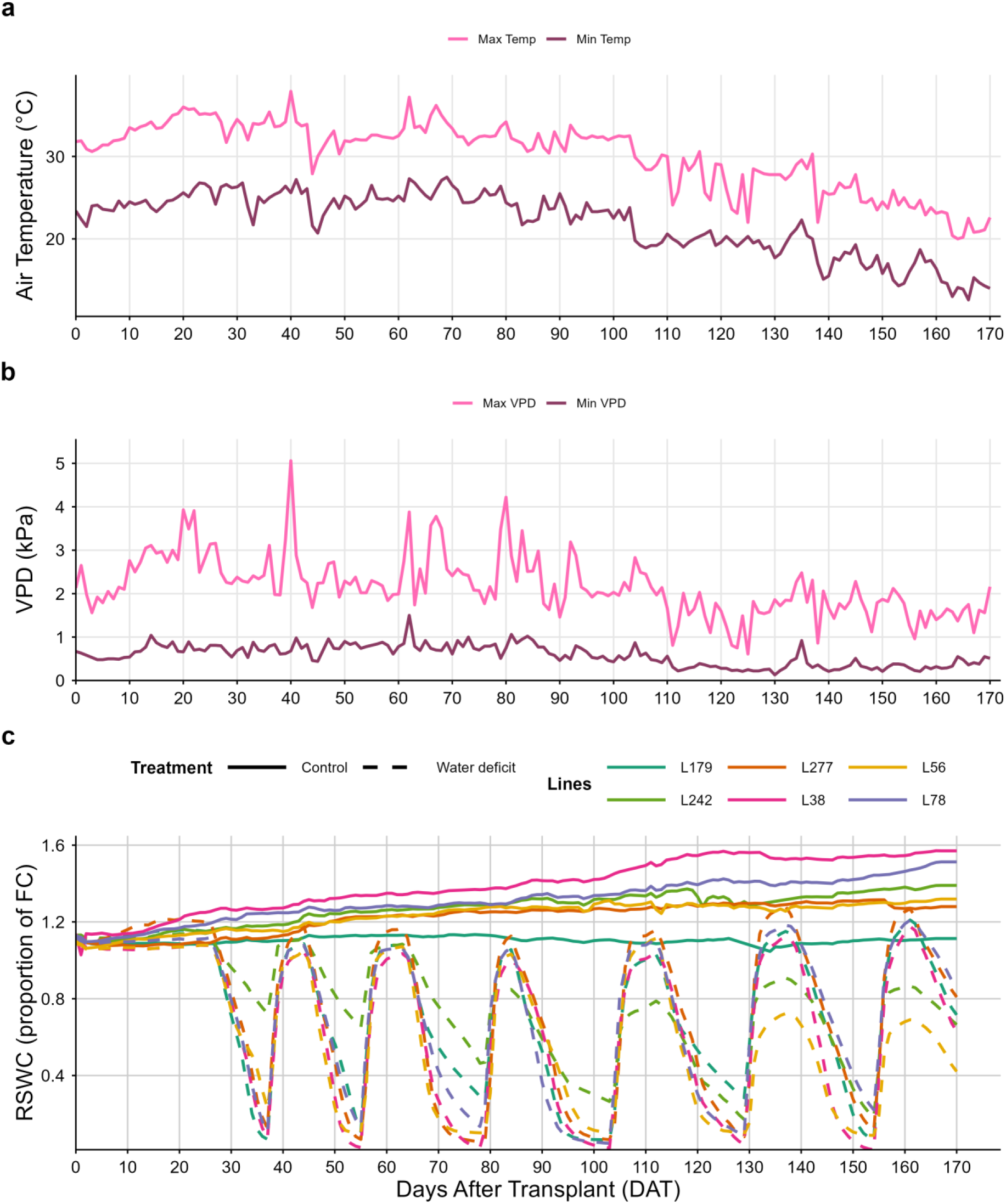
Environmental conditions and soil water status during the experimental period. **(a)** Greenhouse air temperature; **(b)** Vapour pressure deficit (VPD); and **(c)** Relative substrate water content (RSWC) in six eggplant lines under control and water stress conditions. Data in Panel C represent continuous measurements from individual representative plants for each line and treatment.

The plants were irrigated using a fertigation system equipped with one 4 L/h drip emitter per pot. Six 6-minute irrigation cycles were applied per day to maintain the plants at field capacity (FC). The substrate water status was monitored using TEROS 10 sensors (METER Group, Pullman, WA, USA), which recorded measurements every ten minutes. To standardise the data, values were expressed as relative substrate water content (RSWC) relative to field capacity (FC = 1.0); therefore, the daily averages presented represent a proportion of FC rather than absolute volumetric content. The control treatment maintained constant irrigation rates throughout the entire experimental period. The water deficit treatment consisted of withholding irrigation until the RSWC approached 0 in the driest sensors. Five irrigation withdrawal cycles were applied during the trial, lasting 11, 11, 16, 20, and 17 days, respectively (Figure 1). Twelve sensors were used, located in replicates of lines L38, L56, L78, L179, L242, and L277 in both the control and water deficit treatments. Macronutrients were applied via irrigation with two fertiliser solutions (11-3-6 and 7-3-10 N-P-K) at a rate of 0.5 L per 1000 L of irrigation water (Codisa-Dicaher S.L., Valencia, Spain). Furthermore, a solution of Welgro Hydroponic Chelates (Massó, Barcelona, Spain) was applied at a rate of 0.25 L per 1000 L of irrigation water. The composition of the chelates solution was Fe (DTPA) 2.4%, Mn (EDTA) 1.5%, Cu (EDTA) 0.14%, Zn (EDTA) 1.0%, B 0.52%, and Mo 0.12%. Finally, 500 g of humic acid fertiliser (Ascenza, Lisbon, Portugal) was applied per 1,000 L of irrigation water.

### 2.3. Growth and Yield Evaluation

After 170 days after transplanting (DAT), the stem diameter was measured and the fresh biomass of the leaves and stems was weighed to obtain the fresh weight (FW). To calculate the yield, commercially ripe fruits were harvested throughout the trial. The yield under control conditions and the yield under water stress were used to determine the stress tolerance index (STI) [1] (Fernandez, 1992).

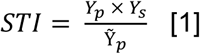

Where Y_s_ is the yield of the line under stress, Y_p_ is the yield of the line under control conditions, and Ỹ_p_ is the average yield of all lines under control conditions.

### 2.4. Physiological analyses

Stomatal conductance (g_s_) was measured at 104 DAT using an SC-1 Leaf Porometer (Meter Group, Pullman, WA, USA), on a fully developed leaf representative of the plant’s water status. Relative water content (RWC) was measured at 104,129, and 154 DAT on representative leaf fragments of approximately 8 cm². These measurements are hereafter referred to as RWC_104_, RWC_129,_ and RWC_154_. The fragment was weighed immediately after cutting to obtain the FW. The fragment was then placed in a 50 ml Falcon tube containing distilled water. After 24 hours, it was weighed again to obtain the turgor weight (TW). Finally, the fragment was placed in an oven at 75 °C for 72 hours to obtain the dry weight (DW), after which the RWC was calculated using the following formula:

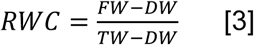

### 2.5. Biochemical analyses

Biochemical analyses were conducted 104 days after transplanting. For all extractions, 0.1 g of fresh leaf tissue was collected from a fully expanded leaf located in the middle of the plant canopy (which was considered to be representative of the plant’s water status). This tissue was then immediately frozen in liquid nitrogen and mechanically homogenised using a beater mixer mill containing glass beads. To avoid the effects of dilution caused by differences in tissue water content among the various treatments, all quantified compounds were expressed on a DW basis.

To quantify the chlorophyll and carotenoid content of each sample, the pigments were extracted from the leaf material using 1 mL of 80% acetone. The samples were kept under agitation in the dark at 4 °C for 24 hours. They were then centrifuged at 13,000 × *g* for 10 minutes at 4 °C to collect the supernatant. Spectrophotometry was then used to measure the samples at 470, 646, and 663 nm. The concentrations of chlorophyll a (Chl a), chlorophyll b (Chl b), and carotenoids (Caro) were obtained using the equations proposed by Lichtenthaler & Wellburn (1983).

The proline content was determined using the Bates et al. (1973) protocol. The extract was obtained by extracting the plant material with 1 mL of 3% (w/v) sulphosalicylic acid. After centrifugation at 13,000 × *g* for 10 minutes at 4 °C, 0.5 mL of the resulting supernatant was collected and mixed with 0.5 mL of acid ninhydrin acid and 0.5 mL of glacial acetic acid. The samples were then heated to 96°C for 60 minutes to allow the reaction to occur. The samples were cooled on ice, 3 mL of toluene was added to extract the proline, and the absorbance was measured at 520 nm. Quantification was performed using a standard curve of L-proline.

The levels of total soluble sugar (TSS), malondialdehyde (MDA), total phenolic compounds (TPC) and total flavonoids (TF) were determined using the same plant extract. Two millilitres of 80% methanol were mixed with the plant material and left to stand in the dark at 4 °C for 24 hours. The mixture was then centrifuged at 13,000 × *g* for 10 minutes at 4 °C to collect the supernatant. TSS quantification was performed using the phenol–sulphuric acid method (Dubois et al., 1956), involving the addition of 0.5 mL of 5% phenol and 2.5 mL of H₂SO₄ to 0.075 mL of the extract. After 20 minutes at room temperature, the absorbance was measured at 490 nm and the concentration determined using a glucose curve.

To quantify MDA, 0.2 mL of the methanol extract was diluted in 0.4 mL of 80% methanol, then mixed with 0.6 mL of 0.5% thiobarbituric acid (TBA) prepared in 20% trichloroacetic acid (TCA). For each sample’s blank, the same volume of extract was mixed with 0.6 mL of 20% TCA. The samples were incubated at 95 °C for 15 minutes and then placed on ice to stop the reaction. Absorbance was measured at 440, 532 and 600 nm, and the equations of Hodges et al. (1999) were used to calculate the MDA concentration. To quantify TPC, 0.1 mL of the methanol extract was diluted in 1.4 mL of water, to which 0.1 mL of the Folin-Ciocalteu regent was added (Blainski et al., 2013). The mixture was left to react for 5 minutes, after which 0.35 mL of 15% Na₂CO₃ was added. The mixture was then left to stand in the dark for 60 minutes. The absorbance of each sample was measured at 765 nm and the concentration calculated in relation to a gallic acid standard curve. TF quantification was carried out according to the method of Zhishen et al. (1999). 0.1 mL of the methanol extract, diluted fivefold in 0.4 mL of water, was used. After five minutes, 0.03 mL of 5% NaNO₂ was added. After a further six minutes, 0.3 mL of 10% AlCl₃ was added, followed by 0.2 mL of NaOH 1M. Absorbance was measured at 510 nm, and the concentration was quantified using a catechin standard curve.

### 2.6. Statistical analysis

The data were processed and analysed using R software (version 4.5.2) (R Core Team, 2025). To evaluate the effects of lines and water treatments, a two-way analysis of variance (ANOVA) was performed using a randomised complete block design (RCBD). The model included line (L), treatment (T), the L×T interaction, and block as fixed effects. The relative importance of each source of variation was determined by its percentage contribution to the total sum of squares (SS). Estimated marginal means (EMM) were calculated using *emmeans* package (Lenth et al., 2025). Mean separations were performed using Fisher’s least significant difference (LSD) test with a significance level of α=0.05. To evaluate genotypic performance under specific conditions, lines were compared within each treatment level. Additionally, the main effect of treatment was analysed independently to determine the overall impact of water deficit on all lines. A principal component analysis (PCA) was performed using the EMMs that had been calculated, using the *FactoMineR* package (Lê et al., 2008). All visualisations were generated with the ggplot2 package (Wickham, 2016).

## 3. Results

### 3.1. Analyses of Variance

A two-way analysis of variance revealed a significant effect of lines and treatments on most traits (p<0.05), except for RWC_104_, g_s_ and TF for lines, and Chl b for treatments (Table 1). The line × treatment interaction had a significant effect on all growth traits (Leaf FW, Stem FW, Stem diameter, Fruit weight, N° Fruits and Yield), water content (RWC_129_ and RWC_154_) and TSS. The line effect was the main contributor to the sum of squares (SS) for most growth traits (Leaf FW, Stem FW, Stem diameter, Fruit weight and N° Fruits), whereas the treatment effect was the main contributor to the SS for physiological traits (RWC_129_, RWC_154_ and g_s_) and Yield. For RWC_104_ and the biochemical traits (Chl a, Chl b, Caro, Proline, TSS, MDA, TPC and TF), the main contributor to SS was the residual effect. The line × treatment interaction was not the main contributor to SS for any trait (Table 1).

**Table 1.** Analysis of variance for the twelve lines under control and water deficit conditions (Treatment), and their interactions (Line × Treatment), across the five blocks.

| Trait | Line | Treatment | Line × Treatment | Block | Residuals |
| --- | --- | --- | --- | --- | --- |
| Leaf FW | 31.3 *** | 22.3 *** | 16.5 *** | 0.8 ns | 29.1 |
| Stem FW | 52.2 *** | 19.1 *** | 15.5 *** | 1.1 ns | 12.2 |
| Stem diameter | 47.3 *** | 18.5 *** | 8.9 ** | 2.8 * | 22.4 |
| Fruit weight | 45.4 *** | 24.1 *** | 6.1 * | 0.4 ns | 24.1 |
| N° Fruits | 35.8 *** | 29.3 *** | 10.5 *** | 0.6 ns | 23.7 |
| Yield | 27.1 *** | 38.2 *** | 10.8 *** | 1.0 ns | 22.9 |
| $RWC_{104}$ | 9.7 ns | 30.2 *** | 7.4 ns | 9.7 ** | 42.9 |
| $RWC_{129}$ | 9.5 * | 39.6 *** | 13.2 ** | 2.7 ns | 35.0 |
| $RWC_{154}$ | 14.2 *** | 35.8 *** | 17.2 *** | 1.9 ns | 30.8 |
| $g_s$ | 4.3 ns | 63.8 *** | 3.0 ns | 0.3 ns | 28.6 |
| Chl a | 16.6 * | 20.5 *** | 4.5 ns | 3.4 ns | 55.1 |
| Chl b | 18.8 * | 1.2 ns | 6.8 ns | 4.4 ns | 68.9 |
| Caro | 18.6 * | 7.8 ** | 5.9 ns | 3.6 ns | 64.2 |
| Proline | 21.1 *** | 28.6 *** | 7.1 ns | 7.8 ** | 35.4 |
| TSS | 28.6 *** | 7.0 *** | 15.6 ** | 5.6 * | 43.1 |
| MDA | 16.0 * | 3.8 * | 14.5 ns | 2.4 ns | 63.3 |
| TPC | 23.4 *** | 21.4 *** | 2.9 ns | 5.8 * | 46.5 |
| TF | 13.1 ns | 11.3 *** | 8.6 ns | 4.4 ns | 62.7 |

The numbers represent the percentage of the sum of squares. \*, \*\* and \*\*\* indicate significant differences with p-values of $< 0.05$ , $< 0.01$ and $< 0.001$ , respectively. FW: fresh weight; $RWC_{104}$ , $RWC_{129}$ , $RWC_{154}$ : relative water content at 104, 129, and 154 DAT; $g_s$ : stomatal conductance; Chl a: chlorophyll a; Chl b: chlorophyll b; Caro: carotenoids; TSS: total soluble sugars; MDA: malondialdehyde; TPC: total phenolic content; TF: total flavonoids.

### 3.2. Growth, Yield, and Tolerance Index

Water restriction severely limited vegetative growth across all 12 evaluated lines. Significant reductions were recorded in leaf fresh weight (46.6%) and stem fresh weight (43.7%) (Table 2). At an individual level, line L56 stood out for its vegetative vigour, maintaining the highest Stem FW under both control and water-deficit conditions. Regarding leaf development, L56 and L101 were the most robust under control conditions; however, water deficit statistically levelled Leaf FW across all lines. Furthermore, an average decrease of 16.3% was observed in stem diameter, with a specific group (L38, L56, L57 and L234) maintaining significantly greater stem thickness under drought conditions (Table 2).

**Table 2.**
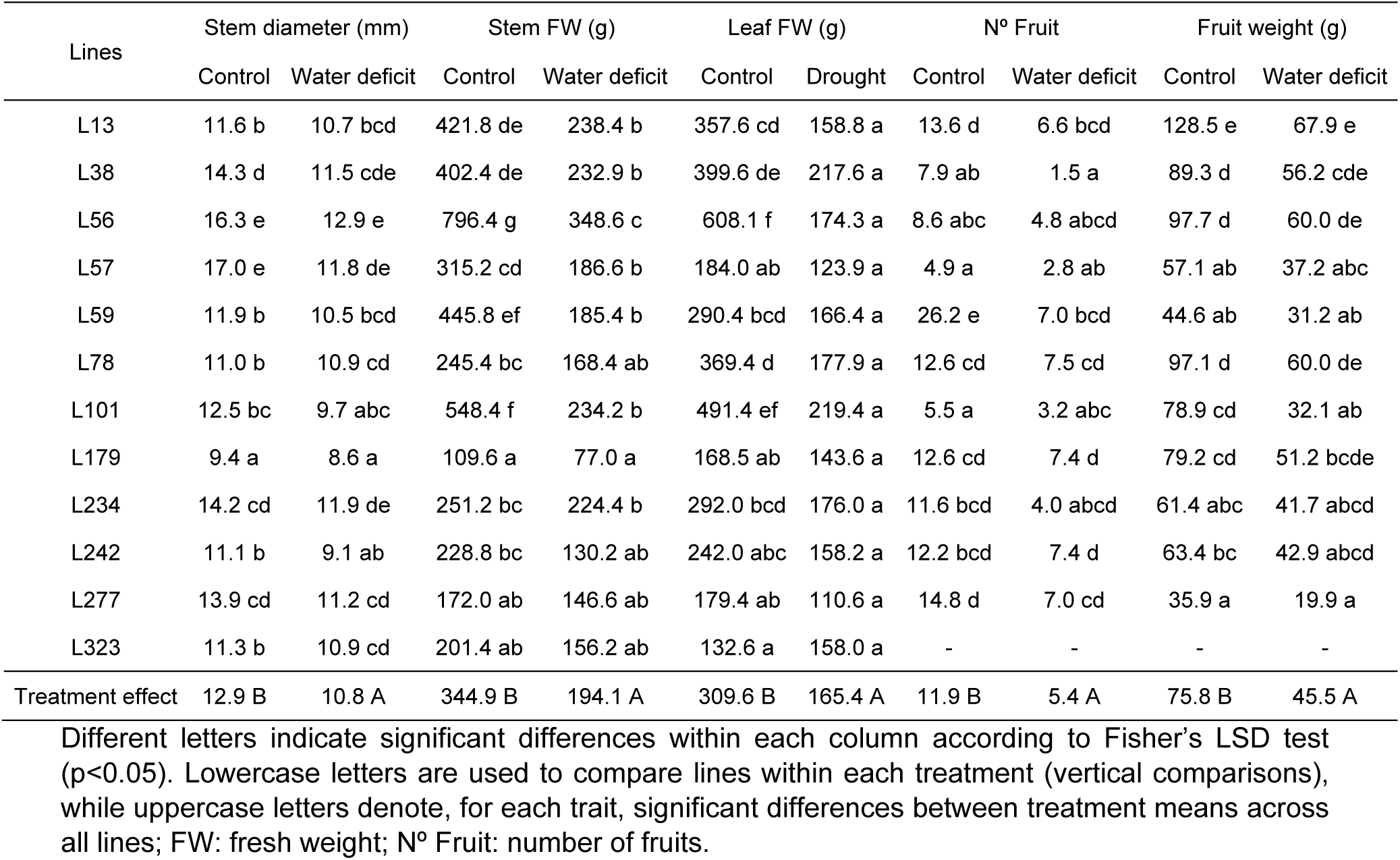
Growth and yield component of the 12 eggplant MAGIC lines evaluated under control and water deficit conditions.

Yield components were impacted more severely than vegetative growth, with average declines of 54.6% in N° Fruit and 40.0% in unit Fruit weight (Table 2). Under water deficit, the ability to sustain yield enabled the identification of resilient genotypes. Eight lines (L13, L56, L59, L78, L179, L234, L242 and L277) were found to significantly outperform the rest, achieving the highest values for N° Fruit. Line L13 showed the highest Fruit weight under both control and water deficit conditions, together with lines L38, L56, L78 and L179. Remarkably, line L323 failed to produce any fruit throughout the trial, regardless of water availability, suggesting a lack of adaptation to the growing environment beyond the hydric factor.

A significant interaction between lines and treatment was observed for yield, indicating contrasting adaptive responses to water deficits. This was evident in substantial shifts in genotype ranking between water regimes (Figure 2a). For example, line L179 showed high plasticity, improving from fourth to second place in terms of average yield under drought conditions. In contrast, other genotypes were more sensitive to stress: L59, for example, was productive under optimal conditions but dropped from third to sixth place when subjected to water deficit. To distinguish between lines with high yield potential and those with robust drought tolerance, the stress tolerance index (STI) was used. This parameter identified a superior cluster of genotypes (L13, L78, and L179) which achieved the highest index scores, highlighting them as the most resilient and productive lines under water deficit (Figure 2b). Conversely, L38, L277, and L57 were identified as the most vulnerable genotypes, recording the lowest STI scores. Notably, the selection based on STI was highly consistent with the previous vegetative stage classification: four of the top five performing lines in the STI ranking had previously been identified as tolerant (L13, L179, L59, and L56).

**Figure 2.**
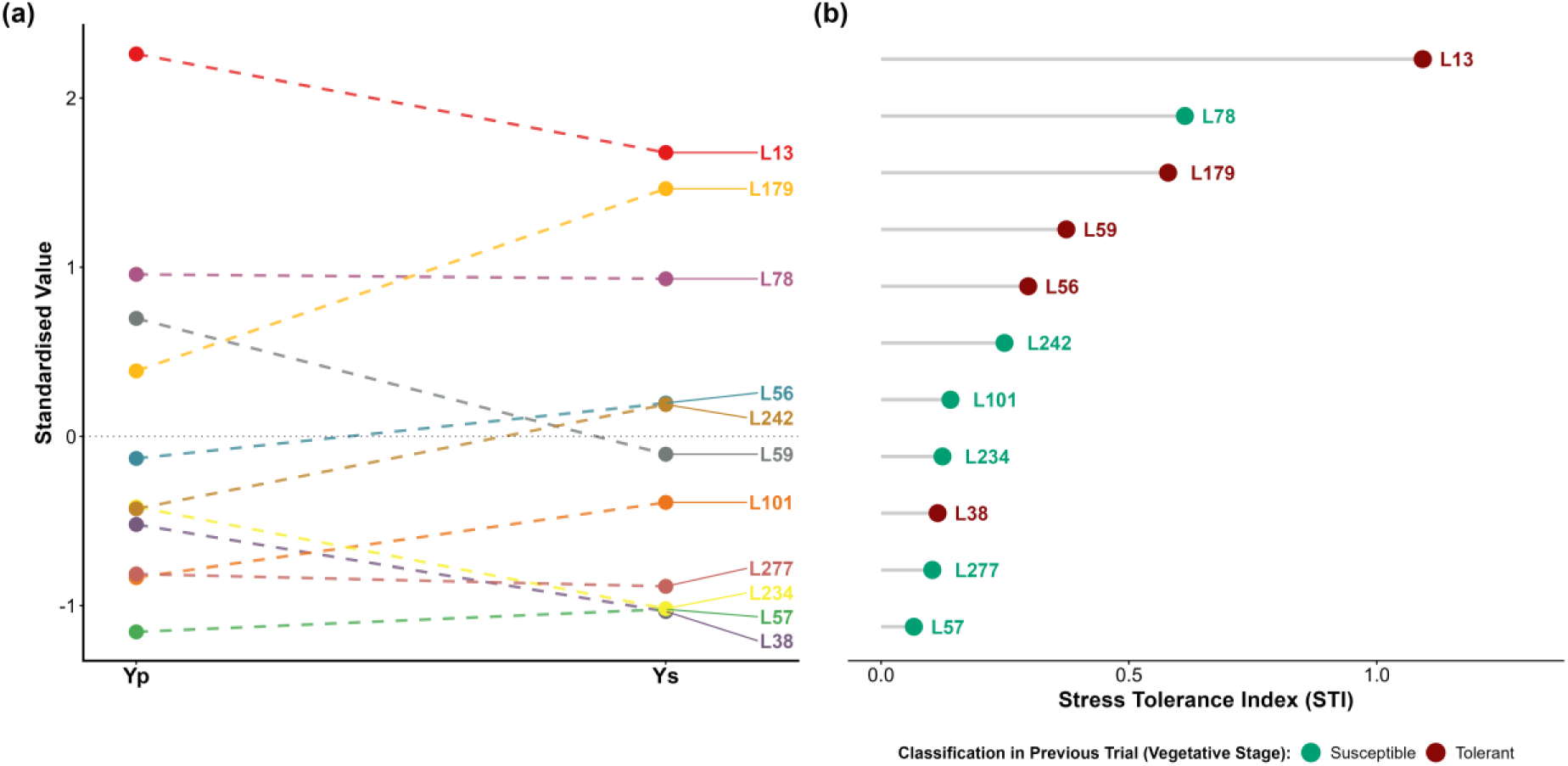
Evaluation and selection of 11 lines based on yield components and the stress tolerance index (STI). (a) Standardized profiles (Z-scores) of yield under control (Yp) and water deficit (Ys) conditions. (b) Ranking of lines ordered from highest to lowest tolerance based on their STI values.

### 3.3. Plant Water Status and Gas Exchange

The water deficit had a significant impact on leaf water status, with relative water content (RWC) decreasing by an average of 23.1%, 25.7% and 21.5% compared to the control at 104, 129 and 154 days after transplanting (DAT), respectively (Table 3). Under control conditions, all lines maintained a consistently high-water status, with values generally exceeding 80.0% and showing no significant differences. In contrast, the lines exhibited distinct physiological responses to water deficit at the three sampling points. At 104 DAT, the highest RWC values were recorded in lines L13, L59, L78 and L277. As the stress progressed to 129 DAT, the top-performing group shifted to include L57, L179, L234, L242 and L323. By the final measurement at 154 DAT, lines L38, L179, L234, L242, L277 and L323 had significantly higher RWC values than the other lines. The lines demonstrated notable shifts in their RWC rankings throughout the trial. For example, line L234 showed clear physiological adaptation, ascending from the lowest RWC group at 104 DAT to become one of the top performers at subsequent time points. In contrast, other lines, such as L56 and L78, exhibited progressive decline in water status (Fig. S1).

**Table 3.** Percentage of leaf relative water content (RWC) in the 12 lines evaluated under control and water deficit conditions at 104, 129 and 154 days after transplant (DAT).

| Lines | 104 DAT |  | 129 DAT |  | 154 DAT |  |
| --- | --- | --- | --- | --- | --- | --- |
|  | Control | Water deficit | Control | Water deficit | Control | Water deficit |
| L13 | 84.2 a | 70.9 bcd | 83.2 a | 50.1 abc | 96.8 a | 63.5 b |
| L38 | 87.8 a | 59.5 ab | 90.8 a | 65.7 cdef | 94.6 a | 79.6 cd |
| L56 | 78.4 a | 63.2 ab | 90.4 a | 58.7 abcd | 90.2 a | 41.7 a |
| L57 | 86.4 a | 56.3 ab | 87.2 a | 73.5 defg | 94.1 a | 70.1 bc |
| L59 | 87.5 a | 87.0 d | 91.9 a | 44.4 a | 93.0 a | 66.7 bc |
| L78 | 81.8 a | 86.2 cd | 91.1 a | 63.2 bcde | 90.5 a | 58.8 b |
| L101 | 83.5 a | 57.2 ab | 83.3 a | 48.8 ab | 85.7 a | 58.3 b |
| L179 | 89.2 a | 64.4 ab | 88.2 a | 78.5 fg | 90.5 a | 87.1 d |
| L234 | 89.6 a | 50.6 a | 83.9 a | 81.5 g | 89.1 a | 79.6 cd |
| L242 | 80.4 a | 62.4 ab | 86.3 a | 74.5 efg | 88.6 a | 77.4 cd |
| L277 | 90.9 a | 69.8 abcd | 94.3 a | 64.9 cdef | 89.4 a | 89.0 d |
| L323 | 90.5 a | 65.8 abc | 82.9 a | 78.4 fg | 90.1 a | 84.9 d |
| Treatment effect | 85.9 B | 66.1 A | 87.8 B | 65.2 A | 91.0 B | 71.4 A |

Different letters indicate significant differences within each column according to Fisher's LSD test ( $p < 0.05$ ). Lowercase letters are used to compare lines within each treatment (vertical comparisons), while uppercase letters denote, for each trait, significant differences between treatment means across all lines.

An average reduction of 67.1% in g_s_ was observed under water deficit conditions compared to control conditions (Figure 3). Under control conditions, significant variability was observed among genotypes, with lines L13, L38, L59, L101, L179, L234, L242 and L323 exhibiting the highest g_s_ values. However, this differentiation disappeared under water stress, where no significant differences were observed between the lines (Figure 3). This indicates a uniform stomatal closure response to drought across the evaluated lines.

**Figure 3.**
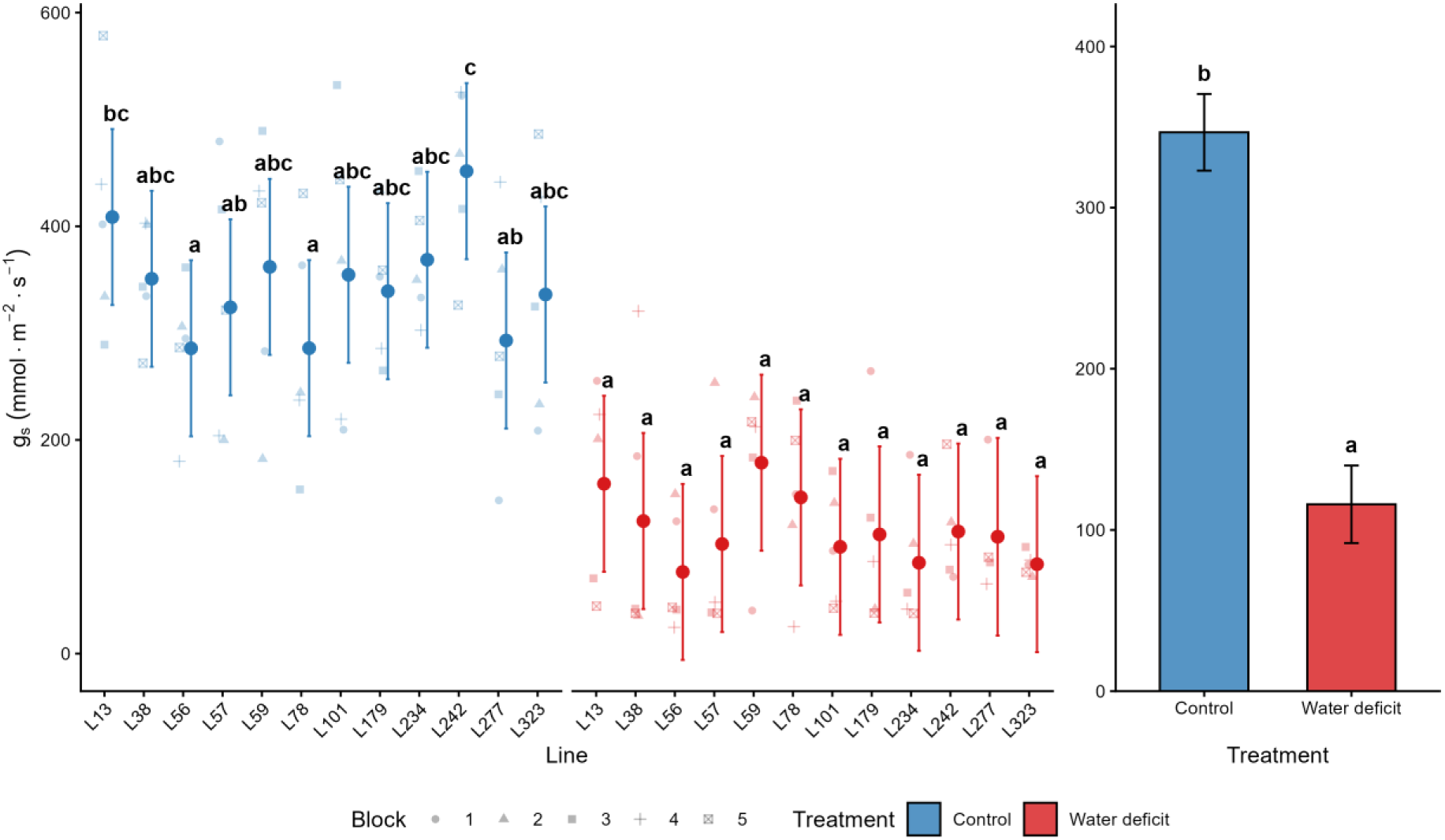
Stomatal conductance (g_s_) in the 12 lines studied under control and water deficit conditions at 104 days after transplant (DAT). Points represent estimated marginal means and error bars indicate 95% confidence intervals. In the line panel (left), different letters indicate significant differences between lines within the same treatment (LSD test; p<0.05). In the global effect panel (right), different letters indicate significant differences between treatment means across all lines (LSD test; p<0.05).

### 3.4. Photosynthetic Pigments and Biochemical Markers

Water deficit caused a significant decline in photosynthetic pigments, with an average reduction of 24.9% in Chl a and 15.2% in Caro. Meanwhile, Chl b levels remained unaffected (Figure 4). Under control conditions, six lines (L59, L78, L101, L179, L242 and L277) exhibited the highest Chl a content. When subjected to water deficit, L59, L101 and L179 were the top performers, significantly outperforming L234, L242 and L323 (Figure 4a). Regarding Chl b, minimal variability was observed under control conditions, with L179 being the only line to significantly exceed L56. However, under water deficit, a broader group of lines maintained higher values: specifically, L38, L56, L59, L78, L101 and L179 (Figure 4b). Finally, under optimal conditions, carotenoid content was highest in lines L38, L57, L59, L78, L101, L179, L242 and L277. Under drought stress, a similar subset of lines, including L38, L56, L59, L78, L101, L179 and L277, maintained this superior performance (Figure 4c).

**Figure 4.**
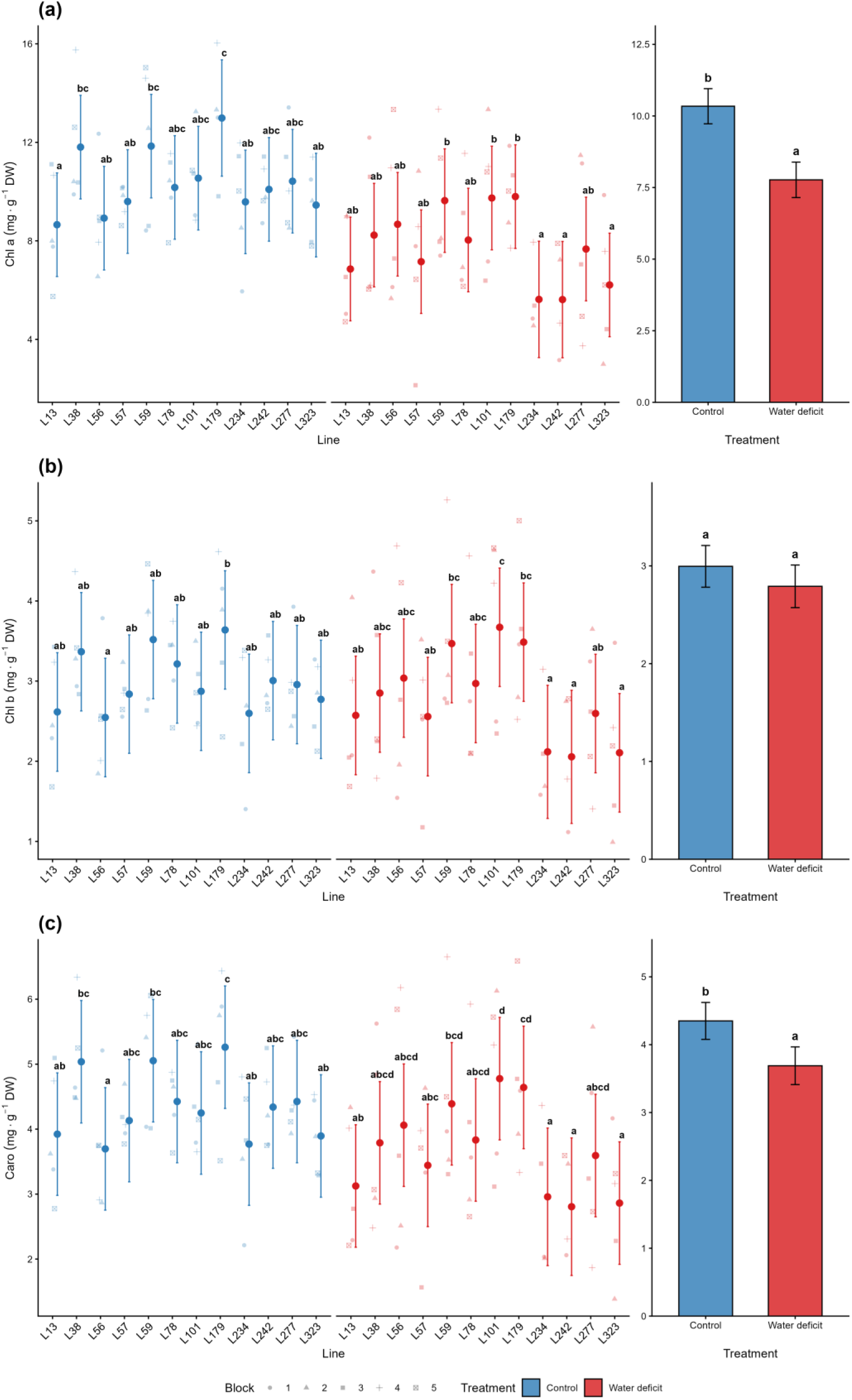
Photosynthetic pigments in the 12 lines studied under control and water deficit conditions at 104 days after transplant (DAT): **(a)** Chlorophyll a (Chl a), **(b)** Chlorophyll b (Chl b) and **(c)** Carotenoids (Caro). Points represent estimated marginal means and error bars indicate 95% confidence intervals. In the line panels (left), different letters indicate significant differences between lines within the same treatment (LSD test; p<0.05). In the global effect panels (right), different letters indicate significant differences between treatment means across all lines (LSD test; p<0.05). DW: dry weight.

A water deficit resulted in the accumulation of osmoprotectants and antioxidant compounds across the evaluated lines. On average, proline content increased by 227.1% under stressful conditions compared to the control. Under optimal irrigation, the highest proline levels were exhibited by lines L56, L101, L242, L277 and L323; however, under water deficit, the most significant accumulation was recorded in line L101, followed by lines L56, L277 and L323 in a superior group (Table 4). Similarly, TSS increased significantly in response to stress (35.4%). While lines L13, L56 and L323 maintained high TSS concentrations under both irrigation regimes, line L323 showed a particularly marked increase under drought conditions, achieving the highest TSS value of all the evaluated lines (Table 4). Regarding oxidative stress markers, MDA increased by an average of 16.6% under water deficit. Under optimal conditions, the highest MDA levels were observed in lines L242 and L101. Under stress conditions, however, line L56 recorded the highest level of oxidative damage. The antioxidant response was characterised by significant increases in TPC (41.9%) and TF (35.3%). Line L323 stood out as the top performer for TPC under both water regimes, particularly under stress. Water availability also influenced the TF response; while lines L242 and L323 led under control conditions, lines L57 and L323 maintained the highest TF content under water deficit (Table 4).

**Table 4.** Proline, total soluble sugar (TSS), malondialdehyde (MDA), total phenolic compound (TPC) and total flavonoids (TF) in the 12 lines evaluated under control and water deficit conditions at 104 days after transplant (DAT).

| Lines | Proline ($\mu\text{mol g}^{-1}\text{ DW}$) | | TSS (mg eq. G $\text{g}^{-1}\text{ DW}$) | | MDA (nmoles $\text{g}^{-1}\text{ DW}$) | | TPC (mg eq. GA $\text{g}^{-1}\text{ DW}$) | | TF (mg eq. C $\text{g}^{-1}\text{ DW}$) | |
| --- | --- | --- | --- | --- | --- | --- | --- | --- | --- | --- |
|  | Control | Water deficit | Control | Water deficit | Control | Water deficit | Control | Water deficit | Control | Water deficit |
| L13 | 4.1 a | 34.9 ab | 26.7 c | 31.7 bc | 348.6 abc | 465.8 bcd | 7.7 ab | 10.0 abc | 1.6 abc | 2.3 ab |
| L38 | 9.2 ab | 70.9 bcdef | 21.2 abc | 20.5 ab | 270.2 a | 402 abcd | 6.8 ab | 10.1 abc | 1.5 abc | 2.0 a |
| L56 | 44.2 c | 79.6 ef | 25.8 bc | 36.4 c | 310.5 ab | 565.6 d | 8.8 bc | 12.2 cd | 2.2 bc | 2.7 ab |
| L57 | 8.1 ab | 42.8 abc | 15.4 abc | 19.2 a | 290.1 a | 343.5 ab | 6.5 ab | 10.8 bc | 1.0 a | 3.0 b |
| L59 | 9.5 ab | 12.6 a | 15.5 abc | 21.9 ab | 440.4 abc | 495.7 bcd | 6.6 ab | 6.6 a | 1.3 ab | 1.8 a |
| L78 | 4.8 ab | 13.7 a | 18.4 abc | 20 ab | 322.9 ab | 493.1 bcd | 6.4 ab | 8.8 ab | 1.5 abc | 2.0 a |
| L101 | 26.8 abc | 91.4 f | 21.3 abc | 17.3 a | 482.8 bc | 353.5 ab | 8.6 abc | 11.1 bc | 2.1 bc | 2.2 ab |
| L179 | 9.8 ab | 53.6 bcde | 20 abc | 21.8 ab | 314.2 ab | 393.1 abcd | 5.3 a | 10.2 abc | 1.5 abc | 2.4 ab |
| L234 | 3.6 a | 69.0 cdef | 15.3 ab | 21.3 ab | 425.4 abc | 277.6 a | 7.4 ab | 10.8 bc | 1.6 abc | 1.9 a |
| L242 | 18.6 abc | 48.9 bcd | 19.2 abc | 23.2 ab | 514.1 c | 449 abcd | 8 abc | 11.2 bc | 2.4 c | 2.3 ab |
| L277 | 34.6 bc | 61.0 bcdef | 12.5 a | 23.6 ab | 279.3 a | 375.4 abc | 5.4 a | 9.5 abc | 1.9 abc | 2.1 ab |
| L323 | 25.7 abc | 73.7 def | 26.2 bc | 65.2 d | 421.2 abc | 541.3 cd | 11.3 c | 15.0 d | 2.3 c | 3.0 b |
| Treatment effect | 16.6 A | 54.3 B | 19.8 A | 26.8 B | 368.3 A | 429.6 B | 7.4 A | 10.5 B | 1.7 A | 2.3 B |

Different letters indicate significant differences within each column according to Fisher's LSD test ( $p < 0.05$ ). Lowercase letters are used to compare lines within each treatment (vertical comparisons), while uppercase letters denote, for each trait, significant differences between treatment means across all lines. DW: dry weight.

### 3.5. Multivariate Analysis

The first two components of the Principal Component Analysis (PCA) accounted for 51.8% and 52.9% of the observed variance under control and water deficit conditions, respectively. For the control group, PC1 and PC2 explained 33.0% and 18.8% of the total variation, respectively (Figure 5a). Under control conditions, lines L59, L78, and L179 clustered in the negative PC1/positive PC2 quadrant, associated with the traits Chl a, Chl b, Caro, N° Fruit, RWC_154_, and Yield. Conversely, L13, L234, L242, and L323 were projected in the positive PC1/PC2 quadrant, close to the vectors corresponding to Fruit weight, g_s_, TSS, MDA, TPC, and TF. Lines L38, L57, and L277 were projected in the negative PC1/PC2 quadrant, linked to RWC_104_ and RWC_129_. Finally, lines L56 and L101 were situated in the positive PC1/negative PC2 quadrant, grouping with Leaf FW, Stem FW, Stem diameter, and Proline. For the water deficit group, PC1 and PC2 accounted for 31.8% and 21.1% of the variation, respectively (Figure 5b). Under stress, the genotype-trait associations shifted: L242, and L323 clustered in the negative PC1/positive PC2 quadrant, driven primarily by RWC_129_, RWC_154_, TSS, TPC, and TF. Notably, the most resilient lines in terms of productivity (L13, L78, and L179) were projected in the positive PC1/PC2 quadrant, showing a strong association with yield-related traits (Fruit weight, N° Fruit, Yield and STI) as well as RWC_104_, g_s_, and MDA. Meanwhile, L38, L56, L57, L101, L234, and L277 grouped in the negative PC1/PC2 quadrant, associated with Stem FW, Stem diameter, and Proline. Lastly, line L59 was located in the positive PC1/negative PC2 quadrant, close to the vectors corresponding to Leaf FW and pigments (Chl a, Chl b, and Caro).

**Figure 5.**
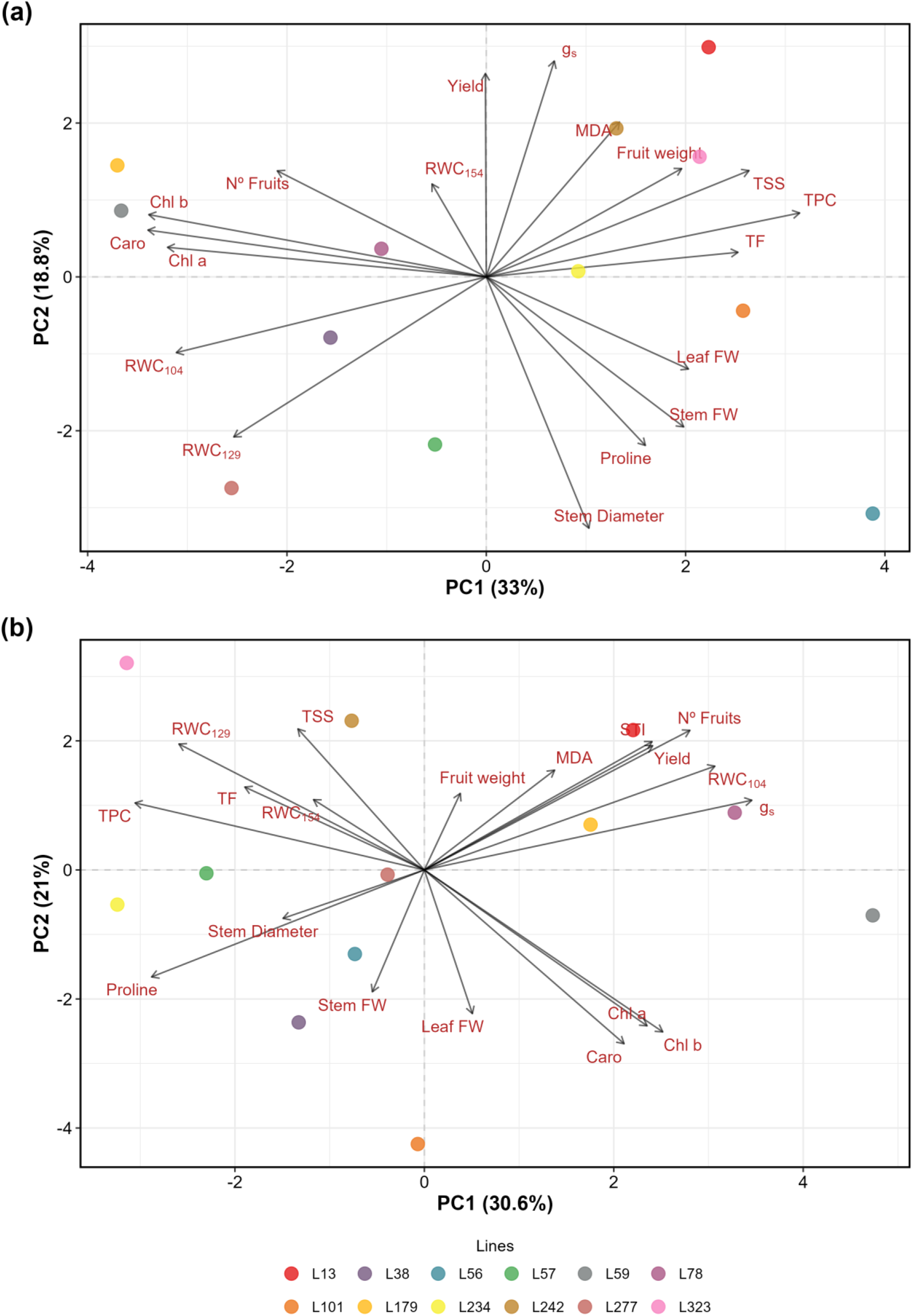
Principal component analysis (PCA) biplot showing the distribution of the 12 eggplant MAGIC lines and traits evaluated under control **(a)** and water deficit **(b)** conditions. FW: fresh weight; RWC_104_, RWC_129_, RWC_154_: relative water content at 104, 129, and 154 d after transplant (DAT); g_s_: stomatal conductance; Chl a: chlorophyll a; Chl b: chlorophyll b; Caro: carotenoids; TSS: total soluble sugars; MDA: malondialdehyde; TPC: total phenolic content; TF: total flavonoids.

## 4. Discussion

Limited water availability is one of the greatest constraints on agricultural production, and breeding drought-tolerant varieties remains one of the most challenging tasks due to the polygenic nature of this trait (Blum, 2011a; Rosero et al., 2020). In our trial, water deficit conditions significantly reduced stem and leaf biomass, directly affecting yield. Integrating the individual yield under stress with the STI successfully identified L13, L78, and L179 as the most resilient genotypes, achieving the highest values and ranking as the three most productive lines under water-deficient conditions.

A key challenge in drought breeding is determining whether tolerance identified in early developmental stages is maintained until maturity. The response to drought has been observed to be stage-specific, depending on the intensity, duration and timing of the stress (Antar et al., 2025). Comparing these results with previous evaluations of the same MAGIC population at the vegetative stage (Flores-Saavedra et al., 2026) revealed a consistent resilience pattern; specifically, lines L13 and L179 maintained their status as top-performing genotypes throughout the entire plant life cycle. This consistency across life stages suggests the presence of stable tolerance mechanisms in these lines, making them excellent candidates for breeding programmes. However, L78’s case highlights the complexity of the responses to water deficit. Although L78 was classified as susceptible during the vegetative phase, it emerged as one of the highest-yielding lines under water stress in this trial. This lack of correlation in some genotypes suggests that tolerance mechanisms during seedling establishment differ from those required for yield maintenance. While screening at the vegetative stage is an effective pre-selection tool, successfully identifying four of our five most tolerant lines (L13, L179, L56 and L59), evaluation at the adult stage is also necessary to identify genotypes such as L78, whose tolerance only manifests during reproductive development. Ideally, a breeding strategy should integrate early-stage screening to identify rapid stress responses and late-stage evaluations to confirm final yield performance, ensuring that all types of genetic tolerance are captured (Araus et al., 2012). Furthermore, these discrepancies may also be attributed to the different methods used to impose water deficit in each study. Tolerance to water stress can vary considerably depending on the environment (Tardieu et al., 2018). In our case study, the vegetative stage assessment maintained a constant field capacity level (Flores-Saavedra et al., 2026); however, the current trial involved irrigation intervals, which subjected adult plants to cyclical periods of stress and recovery. This suggests that lines such as L38 and L78, which exhibit different levels of tolerance in the two trials, may be better adapted to specific stress conditions. In this sense, lines that are tolerant in both conditions are highly valuable as they perform well under different water stress conditions.

Regarding morphological characteristics that may explain production losses, yield components such as the number and weight of fruits decreased significantly under stress. However, the number of fruits was most affected. Other authors have also stated that the number of fruits is the yield component that has the greatest influence on eggplant production under water deficit conditions (Díaz-Pérez and Eaton, 2015; Maachi et al., 2025). Regarding aboveground growth, the most drought-tolerant lines did not exhibit superior growth compared to the others. In fact, line L179, which had the second highest yield under water-deficit conditions, showed low leaf biomass and the least stem growth. This is in contrast to other studies on eggplants which indicate that stem biomass is related to higher productivity (Delfin et al., 2021; Maachi et al., 2025). Moderate leaf growth can reduce transpiration losses and is favoured by good water status under water deficit conditions (Blum, 2005), as was the case with the three drought-tolerant lines.

Of the assessed traits, RWC and g_s_ had the greatest influence on yield under stress conditions and were grouped with yield traits. The most tolerant lines were characterised by high values for these traits under both control and water-deficient conditions. It has been observed in eggplants that higher leaf water content promotes plant growth under water-deficient conditions (Semida et al., 2021). Our findings revealed that plants with greater biomass had lower leaf water content, as larger plants need more irrigation to maintain tissue hydration (Blum, 2011b). Consequently, in the evaluated lines, the water-saving strategy of maintaining lower biomass with optimal water content is beneficial to yield. Furthermore, higher leaf water content was also associated with higher g_s_, as a reduction in water potential (dependent on leaf water status) induces stomatal closure (Chai et al., 2015). In eggplants, g_s_ has been observed to decrease proportionally with water availability (Li et al., 2024a), and the ability to maintain g_s_ levels under stress is considered a tolerance trait (Hannachi et al., 2022; Tani et al., 2018). Although multivariate analysis links g_s_ to higher productivity under water-deficient conditions, no significant differences were observed between the lines.

Photosynthetic pigment maintenance is a key strategy for mitigating abiotic stress (Sharma et al., 2020). Our findings confirm that higher Chl and Caro contents favoured aerial biomass accumulation under water deficit, echoing previous observations in this MAGIC population at the vegetative stage (Flores-Saavedra et al., 2026) and in *S. incanum* introgression lines (Flores-Saavedra et al., 2025). Although a direct correlation between pigments and yield was not universal, the high pigment retention in the top-yielding line L179 highlights this trait as a significant component of its reproductive resilience. Therefore, maintaining these pigments under stress is a fundamental tolerance strategy to preserve photosynthetic capacity (Ashraf & Harris, 2013). This resilience in L179 may be explained by the different susceptibilities of pigments: while Chl *a* within reaction centres is highly prone to oxidative damage, Chl *b* remains structurally stabilized within light-harvesting complexes (LHCII) (Li et al., 2024b). However, other studies in eggplant have observed the degradation of both Chl a and Chl b, although to a greater extent in Chl a (Hannachi et al., 2022; Plazas et al., 2019).

Although proline, an amino acid involved in signalling, antioxidant defences and osmotic adjustment (Ghosh et al., 2022), plays an important role in plant growth under stress, it was not particularly high in the tolerant genotypes. While other studies on eggplants have found that proline enables plants to acclimatise better to stress (Kıran & Furtana, 2023; Sarker et al., 2005), in our trial it appeared to be associated with a higher level of stress. These findings are consistent with previous evaluations of the parents of the MAGIC population (Flores-Saavedra et al., 2024), in which proline accumulation also failed to confer a measurable advantage under water-limiting conditions. In contrast, soil moisture content (SMC) proved to be a more consistent indicator of resilience in the evaluated lines, showing a positive association with maintaining water status and yield. Soluble sugars can contribute to osmotic adjustment and the phytohormone response, thus conferring greater tolerance to plants under water-deficit conditions (Kaur et al., 2021). Our results demonstrate an increase in TSS levels induced by water stress. In the PCA, TSS levels clustered with RWC, and in L13, one of the most tolerant varieties, the TSS content was particularly high. Results from other trials also emphasise the significance of sugars in imparting drought tolerance to eggplants (Chakhchar et al., 2025; Mibei et al., 2018).

Water deficit conditions can induce oxidative stress in plants (Laxa et al., 2019). MDA is a by-product of fatty acid oxidation and can be used to identify the effect of stress on lipid oxidation (Sade et al., 2011). As in other trials investigating the effects of drought stress on eggplants, MDA was found to indicate oxidative stress in certain genotypes (Hannachi et al., 2022; Plazas et al., 2022). Interestingly, multivariate analysis revealed that under both control and water deficit conditions, MDA clustered with the most productive lines. This may be because, although MDA can indicate greater oxidative stress, it also plays a role in acclimatising plants to stress by activating genes involved in response processes that confer greater tolerance (Morales & Munné-Bosch, 2019). In response to oxidative stress, plants can increase the production of certain antioxidant compounds to alleviate stress damage, including TPC, TF and proline (Mishra et al., 2023). These compounds were found to increase in our assay when the plants were subjected to water deficit. Unlike MDA, these three compounds are clearly clustered by PC1, reflecting their coordinated involvement in the antioxidant defense system of the evaluated lines. In line with these results, the antioxidant properties of TPC and TF in eggplants have been consistently observed (Flores-Saavedra et al., 2024; Plazas et al., 2022). While these compounds represent a generalized stress response in eggplant, they do not appear to be the primary drivers of yield stability in this MAGIC population.

## 5. Conclusions

Water deficit has a significant impact on eggplant productivity, primarily by reducing the number and weight of fruits produced. However, the genotypic variability within this MAGIC population enabled lines with contrasting tolerance levels and diverse adaptation strategies to be identified. Our study demonstrates that vegetative-stage selection serves as a powerful predictor of adult-stage performance, as four of the five most tolerant lines identified at the seedling stage maintained their superior performance in the adult trial. This consistency across the entire life cycle confirms that core resilience mechanisms are genetically stable, validating large-scale, cost-effective early screening as an efficient pre-selection tool to identify high-yielding genotypes for water-limited environments. However, as L78 was productive as an adult despite being susceptible as a seedling, it suggests that tolerance mechanisms can be specific to the reproductive stage. This shows that certain genotypes have specific reproductive-stage resilience mechanisms that are not revealed in early-stage stress trials. Drought tolerance in *S. melongena* is clearly a multi-trait characteristic, with resource efficiency (moderate biomass with high hydration) being more beneficial than high vegetative vigour. While L179 excels in pigment preservation, L13 relies on osmotic adjustment through sugar accumulation. Yet both strategies converge on maintaining g_s_ and leaf hydration) as the primary determinants of yield. Finally, while the coordinated accumulation of TPC, TF, and proline reflects a generalised antioxidant response to mitigate oxidative damage, these markers function as indicators of stress intensity rather than primary drivers of yield stability. In summary, integrating early-stage screening with adult-stage validation provides a comprehensive approach for breeding programmes. Lines L13, L179 and L78 represent exceptionally valuable genetic material for developing the next generation of drought-tolerant eggplant.

## CRediT authorship contribution statement

**Martín Flores-Saavedra:** Writing – original draft, Data curation, Formal analysis, Methodology, Investigation. **Mariola Plazas**: Formal analysis, Investigation, Supervision, Writing – review and editing. **Nuria Pascual-Seva:** Conceptualization, Formal analysis, Methodology, Writing – review and editing. **Santiago Vilanova:** Investigation, Supervision. **Oscar Vicente**: Methodology, Supervision, Writing – review and editing. **Pietro Gramazio**: Investigation, Supervision. **Jaime Prohens:** Conceptualization, Formal analysis, Investigation, Supervision, Writing – review and editing.

## Data availability

Data will be made available on request.

## References

Antar, O., Isern, H., Rivera, A., Plazas, M., Díez, M. J., Vilanova, S., & Casals, J., 2025. From greenhouse conditions to the field: Stability of tolerance to water deficit in the tomato wild relatives *Solanum lycopersicum* var. *cerasiforme* and *Solanum pimpinellifolium*. BMC Plant Biology, 25(1), 1682. 10.1186/s12870-025-07582-8

Araus, J. L., Serret, M. D. & Edmeades, G. O., 2012. Phenotyping maize for adaptation to drought. Frontiers in Physiology. 3:305. 10.3389/fphys.2012.00305

Arrones, A., Baraja-Fonseca, V., Solana, A., Plazas, M., Soler, S., Prohens, J., Vilanova, S., & Gramazio, P., 2025. Resequencing and phenotyping of the first highly inbred eggplant multiparent population reveal *SmLBD13* as a key gene associated with root morphology. Horticulture Research, 12(9), uhaf167. 10.1093/hr/uhaf167

Ashraf, M., Harris, & P.J.C., 2013. Photosynthesis under stressful environments: An overview. Photosynthetica, 51, 163–190 (2013). 10.1007/s11099-013-0021-6

Bates, L. S., Waldren, R. P., & Teare, I. D., 1973. Rapid determination of free proline for water-stress studies. Plant and Soil, 39(1), 205–207. 10.1007/BF00018060

Bisbis, M. B., Gruda, N. S., & Blanke, M. M., 2019. Securing horticulture in a changing climate—a mini review. Horticulturae, 5(3). 10.3390/horticulturae5030056

Blainski, A., Lopes, G. C., & Mello, J. C. P. D., 2013. Application and analysis of the Folin Ciocalteu method for the determination of the total phenolic content from *Limonium Brasiliense* L. Molecules, 18(6), 6852–6865. 10.3390/molecules18066852

Blum A., 2005. Drought resistance, water-use efficiency, and yield potential—are they compatible, dissonant, or mutually exclusive? Australian Journal of Agricultural Research, 56, 1159–1168. 10.1071/AR05069

Blum, A., 2011a. Breeding considerations and strategies. In A. Blum (Ed.), Plant breeding for water-limited environments (pp. 235–243). Springer. 10.1007/978-1-4419-7491-4_6

Blum, A., 2011b. Plant water relations, plant stress and plant production. In A. Blum (Ed.), Plant breeding for water-limited environments (pp. 11–52). Springer. 10.1007/978-1-4419-7491-4_2

Chai, Q., Gan, Y., Zhao, C., Xu, H.-L., Waskom, R. M., Niu, Y., & Siddique, K. H. M., 2015. Regulated deficit irrigation for crop production under drought stress. A review. Agronomy for Sustainable Development, 36(1), 3. 10.1007/s13593-015-0338-6

Chakhchar, A., Aitouguinane, M., Afalo, K., Boutabaa, A., Bouhi, R. E., Zidan, M. A., Chabbar, A., & Modafar, C. E., 2025. Integrating morpho-physiological and biochemical traits to assess drought stress and recovery tolerance in eggplant. Notulae Scientia Biologicae, 17(3), 12427–12427. 10.55779/nsb17312427

Çolak, Y. B., Yazar, A., Sesveren, S., & Çolak, İ., 2017. Evaluation of yield and leaf water potantial (LWP) for eggplant under varying irrigation regimes using surface and subsurface drip systems. Scientia Horticulturae, 219, 10–21. 10.1016/j.scienta.2017.02.051

Delfin, E. F., Drobnitch, S. T., & Comas, L. H., 2021. Plant strategies for maximizing growth during water stress and subsequent recovery in *Solanum melongena* L. (eggplant). PLOS ONE, 16(9), e0256342. 10.1371/journal.pone.0256342

Díaz-Pérez, J. C., & Eaton, T. E., 2015. Eggplant (*Solanum melongena* L.) plant growth and fruit yield as affected by drip irrigation rate. HortScience, 50(11), 1709–1714. 10.21273/HORTSCI.50.11.1709

Dubois, Michel., Gilles, K. A., Hamilton, J. K., Rebers, P. A., & Smith, Fred., 1956. Colorimetric method for determination of sugars and related substances. Analytical Chemistry, 28(3), 350–356. 10.1021/ac60111a017

Fernandez, G. C. J., 1992. Effective selection criteria for assessing plant stress tolerance. In Proceedings of the international symposium on adaptation of vegetables and other food crops in temperature and water stress (eds Kuo, C. G.). AVRDC Publication: Tainan, Taiwan: Shanhua: Chapter (25), 257–270. 10.22001/wvc.72511

Ferreira, C. S. S., Soares, P. R., Guilherme, R., Vitali, G., Boulet, A., Harrison, M. T., Malamiri, H., Duarte, A. C., Kalantari, Z., & Ferreira, A. J. D., 2024. Sustainable water management in horticulture: Problems, premises, and promises. Horticulturae, 10(9). 10.3390/horticulturae10090951

Flores-Saavedra, M., Plazas, M., Vilanova, S., Prohens, J., & Gramazio, P., 2023. Induction of water stress in major *Solanum* crops: A review on methodologies and their application for identifying drought tolerant materials. Scientia Horticulturae, 318, 112105. 10.1016/j.scienta.2023.112105

Flores-Saavedra, M., Plazas, M., Gramazio, P., Vicente, O., Vilanova, S., & Prohens, J., 2024. Growth and antioxidant responses to water stress in eggplant MAGIC population parents, F1 hybrids and a subset of recombinant inbred lines. BMC Plant Biology, 24(1), 560. 10.1186/s12870-024-05235-w

Flores-Saavedra, M., Gramazio, P., Vilanova, S., Mircea, D. M., Ruiz-González, M. X., Vicente, Ó., Prohens, J., & Plazas, M., 2025. Introgressed eggplant lines with the wild *Solanum incanum* evaluated under drought stress conditions. Journal of Integrative Agriculture, 24(6), 2203–2216. 10.1016/j.jia.2024.03.014

Flores-Saavedra, M., Bančič, J., Huaman, Y., Arrones, A., Vicente, O., Plazas, M., Vilanova, S., Gramazio, P., & Prohens, J., 2026. Water stress tolerance, genomic selection and identification of genomic regions in a MAGIC population of eggplant. Theoretical and applied Genetic, 139, 153. 10.1007/s00122-026-05251-4

Ghosh, U. K., Islam, M. N., Siddiqui, M. N., Cao, X., & Khan, M. a. R., 2022. Proline, a multifaceted signalling molecule in plant responses to abiotic stress: Understanding the physiological mechanisms. Plant Biology, 24(2), 227–239. 10.1111/plb.13363

Gramazio, P., Alonso, D., Arrones, A., Villanueva, G., Plazas, M., Toppino, L., Barchi, L., Portis, E., Ferrante, P., Lanteri, S., Rotino, G. L., Giuliano, G., Vilanova, S., & Prohens, J., 2023. Conventional and new genetic resources for an eggplant breeding revolution. Journal of Experimental Botany, 74(20), 6285–6305. 10.1093/jxb/erad260

Hannachi, S., Signore, A., Adnan, M., & Mechi, L., 2022. Single and associated effects of drought and heat stresses on physiological, biochemical and antioxidant machinery of four eggplant cultivars. Plants, 11(18). 10.3390/plants11182404

Hodges, D. M., DeLong, J. M., Forney, C. F., & Prange, R. K., 1999. Improving the thiobarbituric acid-reactive-substances assay for estimating lipid peroxidation in plant tissues containing anthocyanin and other interfering compounds. Planta, 207(4), 604–611. 10.1007/s004250050524

Hultgren, A., Carleton, T., Delgado, M., Gergel, D. R., Greenstone, M., Houser, T., Hsiang, S., Jina, A., Kopp, R. E., Malevich, S. B., McCusker, K. E., Mayer, T., Nath, I., Rising, J., Rode, A., & Yuan, J., 2025. Impacts of climate change on global agriculture accounting for adaptation. Nature, 642(8068), 644–652. 10.1038/s41586-025-09085-w

Kaur, H., Manna, M., Thakur, T., Gautam, V., & Salvi, P., 2021. Imperative role of sugar signaling and transport during drought stress responses in plants. Physiologia Plantarum, 171(4), 833–848. 10.1111/ppl.13364

Khalid, M. F., Huda, S., Yong, M., Li, L., Li, L., Chen, Z.-H., & Ahmed, T., 2023. Alleviation of drought and salt stress in vegetables: Crop responses and mitigation strategies. Plant Growth Regulation, 99(2), 177–194. 10.1007/s10725-022-00905-x

Kıran, S., & Furtana, B. G., 2023. Responses of eggplant seedlings to combined effects of drought and salinity stress: effects on photosynthetic pigments and enzymatic and non-enzymatic antioxidants. Gesunde Pflanzen, 75(6), 2579–2590. 10.1007/s10343-023-00901-9

Kumar, R., Solankey, S. S. S., & Singh, M. S., 2012. Breeding for drought tolerance in vegetables. Vegetable Science, 39(01), 1–15.

Laxa, M., Liebthal, M., Telman, W., Chibani, K., & Dietz, K.-J., 2019. The role of the plant antioxidant system in drought tolerance. Antioxidants, 8(4), 94. 10.3390/antiox8040094

Lê, S., Josse, J., & Husson, F., 2008. Factominer: An R package for multivariate analysis. Journal of Statistical Software, 25, 1–18. 10.18637/jss.v025.i01

Lenth, R. V., Piaskowski, J., Banfai, B., Bolker, B., Buerkner, P., Giné-Vázquez, I., Hervé, M., Jung, M., Love, J., Miguez, F., Riebl, H., & Singmann, H., 2025. Emmeans: Estimated Marginal Means, aka Least-Squares Means (Version 2.0.1) [Computer software]. https://cran.r-project.org/web/packages/emmeans/index.html

Li, X., Qiang, X., Yu, Z., Li, S., Sun, Z., He, J., Han, L., Li, Q., & He, L., 2024a. Effects of different water stresses under subsurface infiltration irrigation on eggplant growth and water productivity. Scientia Horticulturae, 337, 113548. 10.1016/j.scienta.2024.113548

Li, X., Zhang, W., Niu, D., & Liu, X., 2024b. Effects of abiotic stress on chlorophyll metabolism. Plant Science, 342, 112030. 10.1016/j.plantsci.2024.112030

Lichtenthaler, H. K., & Wellburn, A. R., 1983. Determinations of total carotenoids and chlorophylls a and b of leaf extracts in different solvents. Biochemical Society Transactions, 11(5), 591–592. 10.1042/bst0110591

Maachi, D., Toppino, L., Ezquer, I., Cebeci, E., Filiz, B. H., Rotino, G. L., & Aberkani, K., 2025. Effect of water supply regimes on physiological parameters and productivity in eggplant grown under mediterranean climate conditions. Plant Biosystems, 159(4), 803–826. 10.1080/11263504.2025.2507635

Mibei, E. K., Owino, W. O., Ambuko, J., Giovannoni, J. J., & Onyango, A. N., 2018. Metabolomic analyses to evaluate the effect of drought stress on selected African Eggplant accessions. Journal of the Science of Food and Agriculture, 98(1), 205–216. 10.1002/jsfa.8458

Mishra, N., Jiang, C., Chen, L., Paul, A., Chatterjee, A., & Shen, G., 2023. Achieving abiotic stress tolerance in plants through antioxidative defense mechanisms. Frontiers in Plant Science, 14. 10.3389/fpls.2023.1110622

Morales, M., & Munné-Bosch, S., 2019. Malondialdehyde: Facts and artifacts. Plant Physiology, 180(3), 1246–1250. 10.1104/pp.19.00405

Muzammal, H., Zaman, M., Safdar, M., Adnan Shahid, M., Sabir, M. K., Khil, A., Raza, A., Faheem, M., Ahmed, J., Sattar, J., Sajid, M., & Zaib, A., 2024. Climate change impacts on water resources and implications for agricultural management. In S. Kanga, S. K. Singh, K. Shevkani, V. Pathak, & B. Sajan (Eds), Transforming agricultural management for a sustainable future: Climate change and machine learning perspectives (pp. 21–45). Springer Nature Switzerland. 10.1007/978-3-031-63430-7_2

Palmgren, M., & Shabala, S., 2024. Adapting crops for climate change: Regaining lost abiotic stress tolerance in crops. Frontiers in Science, 2. 10.3389/fsci.2024.1416023

Plazas, M., González-Orenga, S., Nguyen, H. T., Morar, I. M., Fita, A., Boscaiu, M., Prohens, J., & Vicente, O., 2022. Growth and antioxidant responses triggered by water stress in wild relatives of eggplant. Scientia Horticulturae, 293, 110685. 10.1016/j.scienta.2021.110685

Plazas, M., Nguyen, H. T., González-Orenga, S., Fita, A., Vicente, O., Prohens, J., & Boscaiu, M., 2019. Comparative analysis of the responses to water stress in eggplant (*Solanum melongena*) cultivars. Plant Physiology and Biochemistry, 143, 72–82. 10.1016/j.plaphy.2019.08.031

R Core Team., 2025. R: A language and environment for statistical computing (Version 4.5.2) [Computer software]. https://www.R-project.org/

Ranil, R. H. G., Niran, H. M. L., Plazas, M., Fonseka, R. M., Fonseka, H. H., Vilanova, S., Andújar, I., Gramazio, P., Fita, A., & Prohens, J., 2015. Improving seed germination of the eggplant rootstock *Solanum torvum* by testing multiple factors using an orthogonal array design. Scientia Horticulturae, 193, 174–181. 10.1016/j.scienta.2015.07.030

Rosero, A., Granda, L., Berdugo-Cely, J. A., Šamajová, O., Šamaj, J., & Cerkal, R., 2020. A dual strategy of breeding for drought tolerance and introducing drought-tolerant, underutilized crops into production systems to enhance their resilience to water deficiency. Plants, 9(10). 10.3390/plants9101263

Sade, B., Suuml, Soylu, L., & Yetim, E., 2011. Drought and oxidative stress. African Journal of Biotechnology, 10(54), 11102–11109. 10.5897/AJB11.1564

Sarker, B. C., Hara, M., & Uemura, M., 2005. Proline synthesis, physiological responses and biomass yield of eggplants during and after repetitive soil moisture stress. Scientia Horticulturae, 103(4), 387–402. 10.1016/j.scienta.2004.07.010

Semida, W. M., Abdelkhalik, A., Mohamed, Gamal. F., Abd El-Mageed, T. A., Abd El-Mageed, S. A., Rady, M. M., & Ali, E. F., 2021. Foliar application of zinc oxide nanoparticles promotes drought stress tolerance in eggplant (*Solanum melongena* L.). Plants, 10(2), 421. 10.3390/plants10020421

Sharma, A., Kumar, V., Shahzad, B., Ramakrishnan, M., Singh Sidhu, G. P., Bali, A. S., Handa, N., Kapoor, D., Yadav, P., Khanna, K., Bakshi, P., Rehman, A., Kohli, S. K., Khan, E. A., Parihar, R. D., Yuan, H., Thukral, A. K., Bhardwaj, R., & Zheng, B., 2020. Photosynthetic response of plants under different abiotic stresses: A review. Journal of Plant Growth Regulation, 39(2), 509–531. 10.1007/s00344-019-10018-x

Tani, E., Kizis, D., Markellou, E., Papadakis, I., Tsamadia, D., Leventis, G., Makrogianni, D., & Karapanos, I., 2018. Cultivar-dependent responses of eggplant (*Solanum melongena* L.) to simultaneous *Verticillium dahliae* infection and drought. Frontiers in Plant Science, 9. 10.3389/fpls.2018.01181

Tardieu, F., Simonneau, T., & Muller, B., 2018. The Physiological Basis of Drought Tolerance in Crop Plants: A Scenario-Dependent Probabilistic Approach. Annual Review Plant Biology. 69:733–759. 10.1146/annurev-arplant-042817-040218

Toppino, L., Barchi, L., & Rotino, G.L., 2022. Next Generation Breeding for Abiotic Stress Resistance in Eggplant. In: Kole, C. (eds) Genomic designing for abiotic stress resistant vegetable crops. Springer, Cham. 10.1007/978-3-031-03964-5_4

Wickham, H., 2016. ggplot2: Elegant graphics for data analysis. https://ggplot2.tidyverse.org

Zhishen, J., Mengcheng, T., & Jianming, W., 1999. The determination of flavonoid contents in mulberry and their scavenging effects on superoxide radicals. Food Chemistry, 64(4), 555–559. 10.1016/S0308-8146(98)00102-2

