## Supplementary material for "Impact of Water Deficit on Growth, Biochemical, and Physiological Traits in Eggplant MAGIC Lines": Figure S1

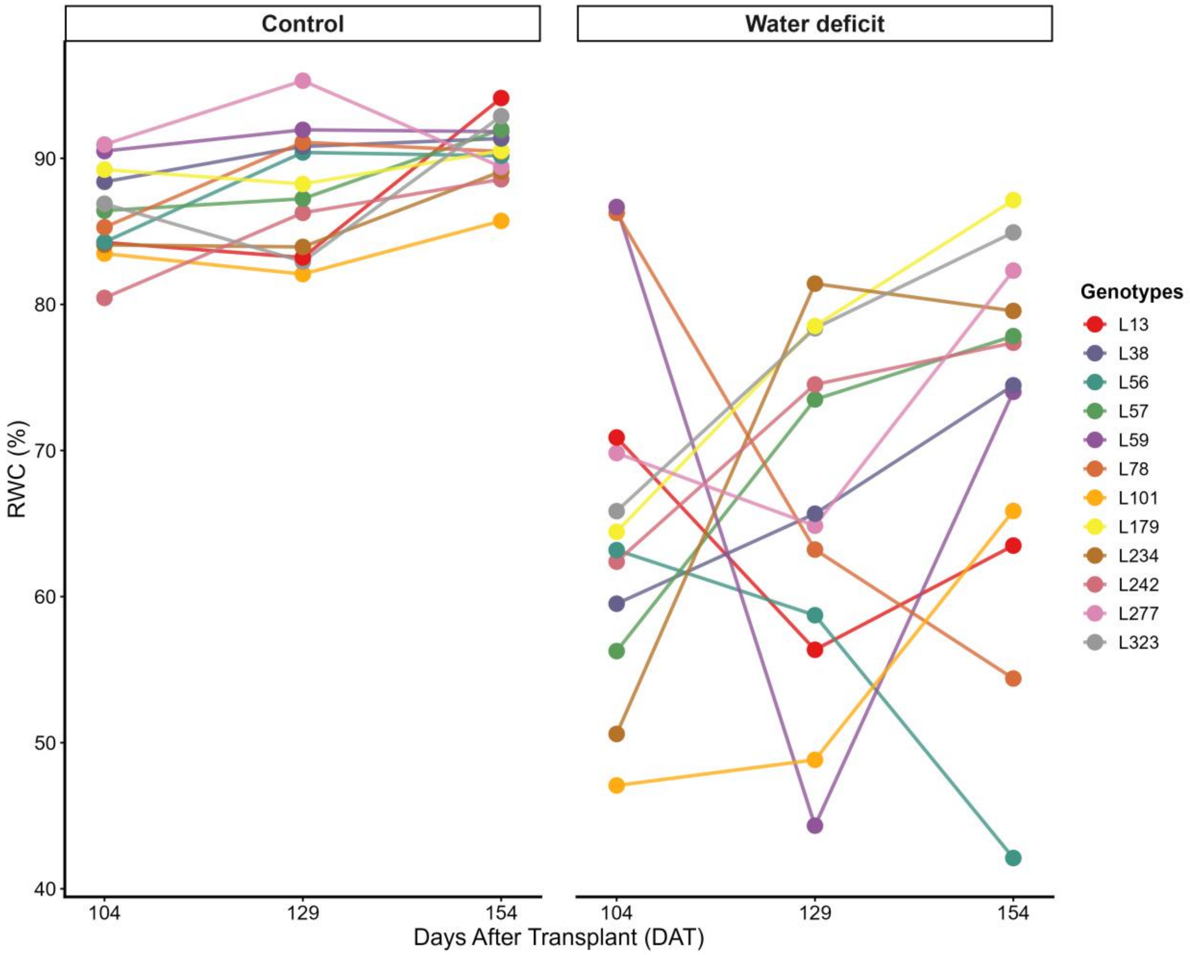


**Fig. S1.** Leaf relative water content (RWC) in the 12 lines studied under control and drought conditions at 104, 129 and 154 days after transplant (DAT).
